# Beyond Turing: *Far-from-equilibrium* patterns and mechano-chemical feedback

**DOI:** 10.1101/2021.03.10.434636

**Authors:** Frits Veerman, Moritz Mercker, Anna Marciniak-Czochra

**Affiliations:** University of Leiden, Mathematical Institute, Niels Bohrweg 1, 2333 CA Leiden, The Netherlands; Heidelberg University, Institute for Applied Mathematics, Im Neuenheimer Feld 205, 69120 Heidelberg, Germany; Heidelberg University, Institute for Applied Mathematics and Interdisciplianry Center for Scientific Computing (IWR), Im Neuenheimer Feld 205, 69120 Heidelberg, Germany

**Keywords:** pattern formation, evolution equations, reaction-diffusion, mechano-chemical models, *far-from-equilibrium* patterns

## Abstract

Turing patterns are commonly understood as specific instabilities of a spatially homogeneous steady state, resulting from activator-inhibitor interaction destabilised by diffusion. We argue that this view is restrictive and its agreement with biological observations is problematic. We present two alternative to the ‘classical’ Turing analysis of patterns. First, we employ the abstract framework of evolution equations to enable the study of *far-from-equilibrium* patterns. Second, we introduce a mechano-chemical model, with the surface on which the pattern forms being dynamic and playing an active role in the pattern formation, effectively replacing the inhibitor. We highlight the advantages of these two alternatives vis-à-vis the ‘classical’ Turing analysis, and give an overview of recent results and future challenges for both approaches.

## 1 Introduction

The prevailing ‘Turing theory’ of pattern formation in biology, and treatment of associated ‘Turing patterns’, can be traced to the work of Hans Meinhardt, germinating from his seminal paper with Alfred Gierer [1]. While Turing in his pioneering paper [2] introduced the framework of reaction-diffusion equations, and the different scenarios in which pattern-like solutions can occur, Gierer and Meinhardt took a distinctly more ‘modelling’ point of view. They introduced the concept of local activation versus long range inhibition, and argued there (and in subsequent work [3, 4, 5]) why this mechanism is not only sufficient, but also necessary, to explain *de novo* pattern formation. This viewpoint has inspired and guided a significant portion of pattern formation research in the last decades [6, 7, 8, 9, 10, 11]. While Turing-type models are based on intercellular signaling through diffusive molecules, they may also arise in continuous-limit of a direct cell-to-cell communication through cell-membrane molecules. Other models of pattern formation are based on different mechanisms of signal transduction such as chemotaxis [12, 13], haptotaxis [14], phase shift in signaling between discrete cells [15] or mechanical forces [16, 17].

To go ‘beyond Turing’, we first briefly discuss pivotal aspects of the canonical ‘Turing’ approach to pattern formation. This discussion will be, by its brevity, both schematic and inherently incomplete. One should be aware of the danger that such a schematic picture can, inadvertently, lead to a caricature description of the theory of Turing patterns, thereby dressing up a straw man that can easily be put aside in favour of new, ‘better’ approaches. With that in mind, our aim is to highlight certain defining viewpoints inherent to the canonical Turing approach, and to present alternatives to these viewpoints, describing situations where these alternatives might be worth considering.

### 1.1 Formation of Turing patterns

Generally speaking, when the terms ‘pattern formation’ and/or ‘Turing patterns’ are used, the following mechanism is assumed to play a role.

1. The system under consideration admits a *stationary, spatially homogeneous steady state*.
2. The temporal evolution of the system is described by certain evolution rules. In the context of these rules, one assumes that the stationary homogeneous steady state is *unstable* for spatially heterogeneous perturbations. This instability is assumed to be of a certain type, so that there exist certain ‘modes’ exhibiting spatial regularity, that are thought to induce a pattern forming process.
3. When initialising the system close to the stationary homogeneous steady state, the early evolution of such an initial state is assumed to be dominantly determined by these ‘pattern modes’, thereby initiating the growth of a ‘pattern’.
4. The spatial heterogeneity of the solution, induced by the pattern modes, is assumed to become more pronounced under the evolution of the system. Eventually, the system settles into a stationary configuration that has the *same* spatial shape as the dominant ‘pattern mode’. This solution is called a ‘Turing pattern’.

To summarise, the necessary ingredients are a stationary, homogeneous steady state, that is unstable in such a way that ‘Turing modes’ dominate the initial evolution – these ingredients then enable the spontaneous formation of a pattern by self-organisation.

### 1.2 Evolution equations

The above description of pattern formation and Turing patterns is deliberately kept vague in terms of the details of the system under consideration, and the ‘rules’ that govern its evolution. This is to highlight the methodology, rather than the specific modelling context – indeed, a pattern forming process as described here, can take place in a wide variety of models including systems of reaction-diffusion equations [18], non-local models of integro-differential equations [19, 20], degenerated reaction-diffusion-ODE systems [21, 22, 23] or fourth order partial differential equation models accounting for evolution of thin elastic structures [17, 24]. Hence, it seems useful to adapt a rather abstract approach, and describe the system as the evolution of a state variable *u*(*x*, *t*) ∈ ℝ^*n*^, determined by the evolution equation

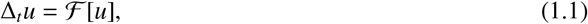

where the functional 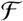 encodes the evolution ‘rules’. In the context proposed by Turing and used by Gierer and Meinhardt, namely that of reaction-diffusion equations, the state variable *u*(*x*, *t*) obeys a system of parabolic partial differential equations in the variables *t* ≥ 0 and *x* ∈ Ω ℝ^*m*^, which can be cast in this abstract framework by choosing

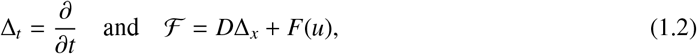

where Δ_*x*_ is the *m*-dimensional Laplacian with, for example, zero-flux boundary conditions, *D* is an *n* × *n* diffusion matrix (which might depend on *u*), and *F* : ℝ^*n*^ → ℝ^*n*^ encodes the ‘reaction terms’. It is important to note that the class of evolution equations (1.1) consists of more than ‘just’ reaction-diffusion systems: Δ_*t*_ can also be chosen to describe a discrete time step, for example Δ_*t*_*u* = *u*(·, *t* + 1) − *u*(·, *t*), where *t* ∈ ℕ; in addition, the spatial variable *x* can be chosen to be discrete, such that *x* ∈ ℤ^*m*^; furthermore, the evolution operator 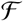 might describe not only deterministic, but also stochastic evolution.

Whatever the context, the abstract formulation (1.1) suggests the viewpoint of dynamical systems theory. That is, for a suitably chosen phase space 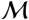, the functional 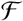 can be seen as a map from 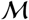 to 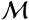, and the evolution equation (1.1) induces a (semi-)flow on this phase space. In this formulation, stationary states are equilibria of this flow, and are therefore equivalent to a solution *u*_∗_ to the equation

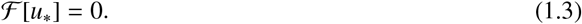

Whether such a stationary state is spatially homogeneous as well, depends on the details of the operator 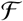 – in particular, the manner in which 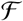 depends on *x*. Note that, in the context of reaction-diffusion equations (1.2), spatially homogeneous stationary states are determined by the reaction terms *F*(*u*) only, as they are solutions to the equation *F*(*u*_∗_) = 0 (1.2).

In this abstract context, formation of Turing patterns as described above, is depicted in Figure 1 A. We would like to highlight two important aspects suggested by this view. First, the flow of the system steers the evolution of an initial state *u*_0_ that is close to the stationary homogeneous state *u*_∗_, towards a second stationary state *u_p_*, which we call the *pattern*, which is not spatially homogeneous, but displays a spatial regularity. Second, this ‘target state’ *u_p_* must be stable in some sense, as it is the ‘endpoint’ of the evolution of a certain initial state *u*_0_. Indeed, if we demand that this pattern formation process does not sensitively depend on the specifics of the initial state *u*_0_ – that is, that an entire open neighbourhood of the stationary homogeneous state *u*_∗_ is attracted to, and flows towards, this stationary pattern-like equilibrium *u_p_* – then the pattern equilibrium *u_p_* must be *stable*, and the stationary homogeneous steady state *u*_0_ must lie in (the interior of) the *basin of attraction* of the equilibrium *u_p_*. In many cases, this is ensured by assuming that the pattern state *u_p_* is *close* to the homogeneous equilibrium *u*_∗_ – note that this implies the need for a notion of distance in phase space, as we have to specify and quantify what we mean by ‘close’. As a result, Turing patterns in reaction-diffusion systems, where the prime candidate for the phase space is the Hilbert space *L*^2^, often have relatively small amplitude.

**Figure 1:**
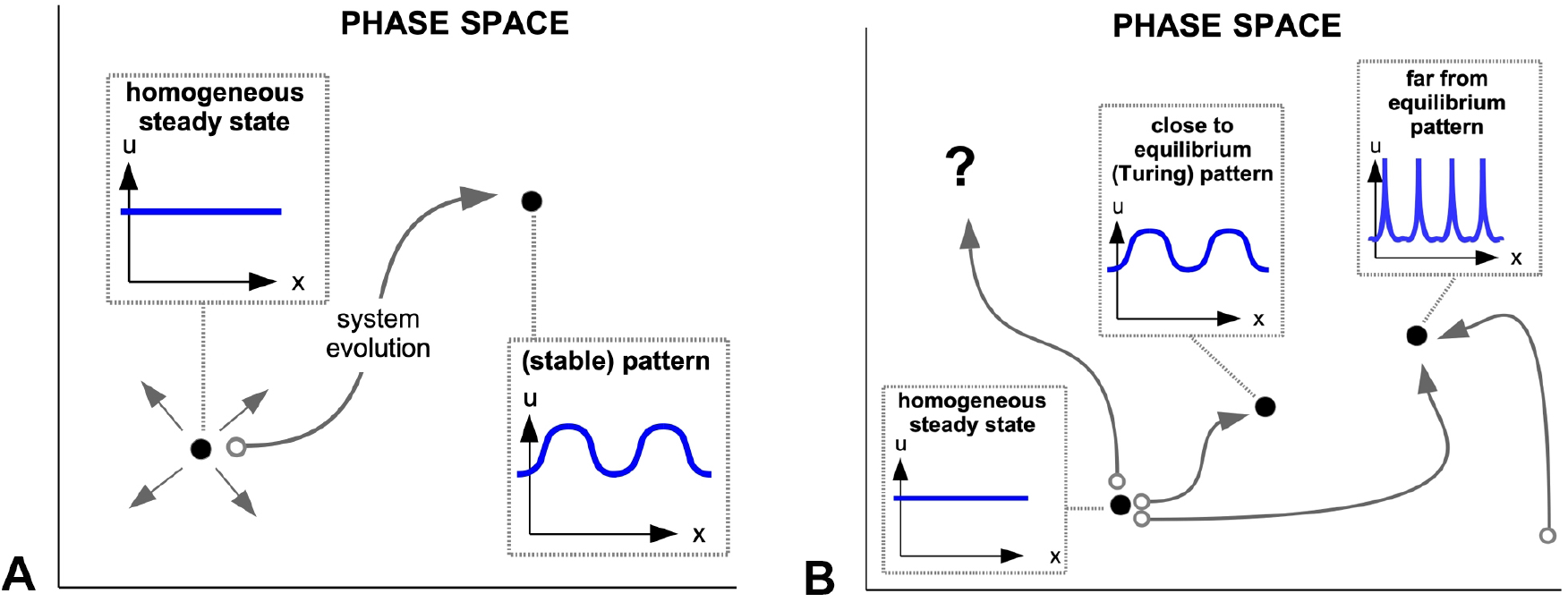
(A) The formation of a Turing pattern in the abstract context of evolution equations. (B) Possible evolution scenarios.

Described in this manner, the conditions for Turing pattern formation seem to be very special indeed. We are invited by abstract viewpoint of evolution equations to ask whether these ‘Turing conditions’ are necessary or sufficient for patterns to occur. In the ‘dynamical systems’ context described above, it seems worthwhile to investigate whether the following scenarios may occur, see also Figure 1 B:

1. An initial state *u*_0_ close to the spatially homogeneous stationary state *u*_∗_ may evolve to an equilibrium that is *far away* from *u*_∗_. If this equilibrium has pattern-like qualities, it is called a *far-from-equilibrium pattern*.
2. An initial state *u*_0_ close to the spatially homogeneous stationary state may *not* evolve towards an equilibrium at all; its evolution might be unbounded, it might evolve towards a chaotic attractor, or towards a non-trivial invariant set in phase space, such as a (time-)periodic orbit or a solution exhibiting mass concentration or a blow-up in finite or infinite time.
3. We may initialise the system *not* close to the spatially homogeneous stationary state *u*_∗_, and this initial state may yet evolve towards a stationary pattern.

The feasibility of these scenarios will be discussed in section 2.

### 1.3 Pattern formation mechanisms

One might object to, or have reservations about, the highly abstract viewpoint of evolution equations, as introduced above. After all, models for specific biological systems do not appear out of thin air, but are carefully constructed based on deep knowledge of, and experience with, the underlying biological phenomena. As such, the building blocks of these models (diffusion or advection processes, specific non-linear reaction terms, etc.) aim to have specific biological meaning, as to provide a faithful description of biological, observed, reality. From a modelling perspective, the evolution equation context does not seem to give any guidance towards the question which terms to include, or which terms to leave out, for the model to exhibit a desired behaviour – such as the occurrence and formation of spatial patterns. Perhaps surprisingly, there turns out to be a general framework in which such specific guidances can be obtained; this will be discussed in section 2. For now, we continue our discussion of the ‘modelling’ perspective.

The local activation / long range inhibition (LALI) approach as advocated by Gierer and Meinhardt has proven to be very effective in producing pattern-like structures, in a range of different model contexts [4]. The efficacy of this approach is directly related to two concepts that, when intertwined in the manner suggested by Gierer and Meinhardt, promote self-organisation and the formation of spatially periodic structures. First, in order to make a meaningful distinction between *local* and *long range*, the biological process (and the model describing that process) needs to incorporate a form of *spatial scale separation*, or multiple spatial scales. Second, there need to be identified two separate model components that interact in a specific way, forming an activator-inhibitor pair. The implementation of this interaction might differ from model to model, but the general idea is that the activator *a* promotes its own growth as well as that of the inhibitor *h*, while the latter inhibits the growth of *a*. When the inhibitor acts on a spatially long range, while the action of the activator is more spatially localised, the emergence of periodic patterns is thought to occur spontaneously; see Figure 2 A.

**Figure 2:**
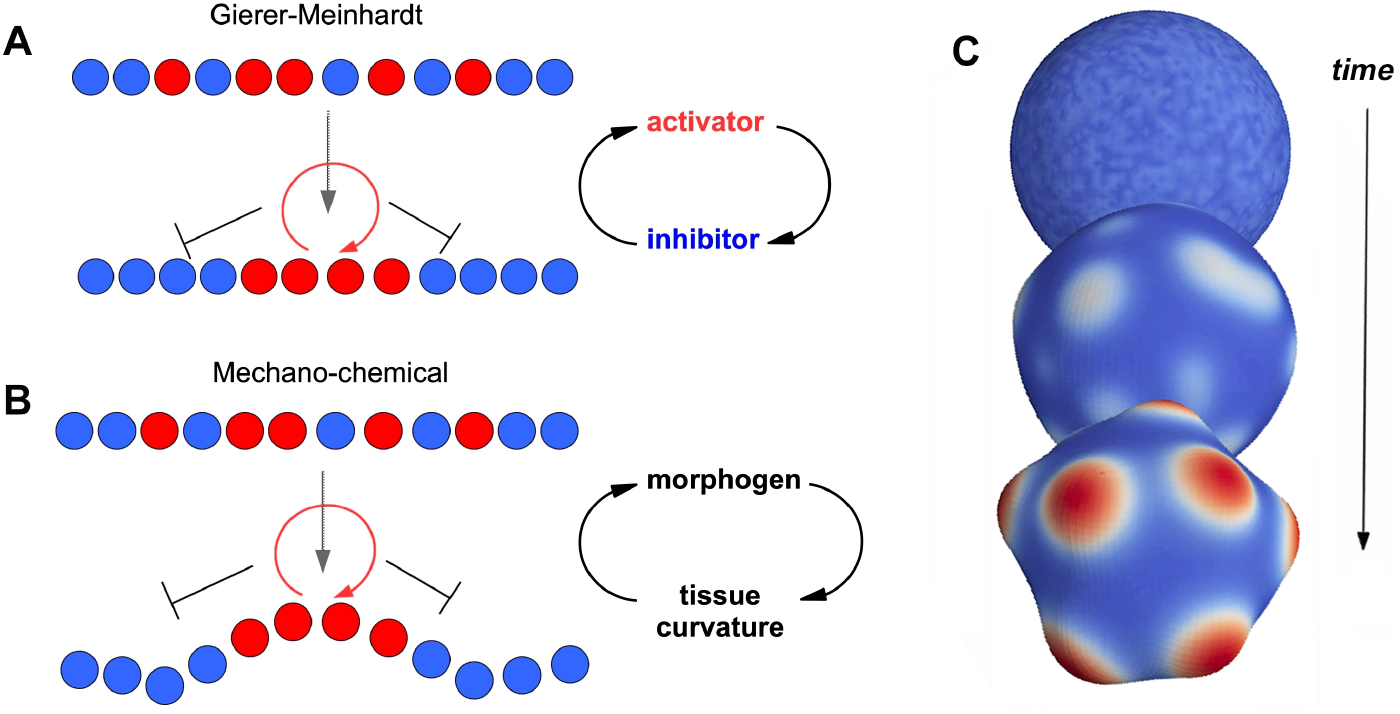
(A) The local activation / long range inhibition idea of Gierer and Meinhardt. (B) Mechanochemical feedback, curvature takes the role of the long range inhibitor. (C) Self-organized pattern formation based on a simple positive feedback loop between morphogen production and local tissue curvature.

In the model context of reaction-diffusion equations, this idea is implemented by studying the interaction of two diffusing species (i.e. *u*(*x*, *t*) = (*a*(*x*, *t*), *h*(*x*, *t*))), where a significant difference in diffusion rates between the two species induces a spatial scale separation in the spatial spread, and therefore the action, of these two species. The specific activator-inhibitor interaction is encoded in the reaction terms *F*(*u*) (1.2).

The success and impact of the LALI approach rests on two pillars: First, the mechanism underlying the formation of patterns is compelling, easy to understand, and sufficiently general to implement in a wide range of biological contexts. Second, when implemented in a specific model context, numerical simulations robustly show the emergence of spatial patterns, the shape of which can be adjusted to match biological observations and experiments by tweaking the model (e.g. [25, 5]). Hence, the approach possesses strong explanatory and predictive qualities, which make it a prime candidate for the explanation of several spatial patterns observed in nature. However, when merging this theory of pattern formation with biological practice, a hurdle presents itself, which prevents the direct application of the ideas of Gierer and Meinhardt. The difficulty lies in the fact that the theory needs (at least) *two* independent, diffusing, species, that in addition must form an activator-inhibitor pair, i.e. interact in the specific manner described above. In very many applications, this poses an insurmountable problem. While a prime candidate for the activator can often be identified without much difficulty, the same is not the case for the inhibitor. Either a long-range (fast diffusing) agent exists, but does not interact with the activator in the required fashion, or an inhibitory agent does not manifest the required spatial long range action (e.g. [26]). In both cases, the proposed inhibitor remains elusive.

Nevertheless, the LALI idea remains singularly persuasive. Its simplicity endows it with a ring of truth: When patterns are observed, the LALI mechanism *must* be its cause, therefore an inhibitor *must* be present. This conviction has guided the experimental efforts in several biological contexts where patterns are observed (e.g. [26, 5]).

We perceive this discrepancy between theory and practice as a strong motivation to go ‘beyond Turing’ in the following sense. We propose to challenge the ‘Turing’ paradigm, implemented in the specific manner advocated by Gierer and Meinhardt, and present an alternative explanatory principle, namely that of *mechano-chemical feedback*. This novel principle has three distinct selling points. First, the simple, convincing aspects of the principle of LALI are preserved, thereby acquiring its persuasive qualities as an explanatory principle. Second, the need for a ‘second species’ that acts as an inhibitor disappears, automatically providing a better potential match with biological practice. Third, the spatial background (i.e. the biological environment) now plays an active and pivotal role, in contrast to the ‘classical Turing’ approach, where the spatial model variables are mostly taken to be in a featureless, Euclidean space ℝ^*m*^ – where, in contrast, biological patterns often form on a dynamic background and thus under continually changing conditions [27]. The principle of mechano-chemical feedback can therefore be viewed as equally compelling, more simple, and closer to biological practice than the ‘classical Turing’ pattern formation principle. The mechano-chemical approach to pattern formation will be discussed in more detail in section 3.

### 1.4 Beyond Turing: Change of perspective, change of paradigm

Both the study of far-from-equilibrium patterns and the introduction of mechano-chemical feedback present ways to go ‘beyond Turing’. The abstract viewpoint of evolution equations, where the study of patterns as the final stage of an evolutionary process, presents a change of perspective. One could describe this as a focal shift from de novo pattern *formation* towards pattern *analysis*. Indeed, for many far-from-equilibrium patterns, or even for more general spatio-temporal ‘dynamic’ patterns such as rotating spirals [28, 29], the question is not so much where the system was initialised, but to which stable states the system can evolve towards. This frees oneself from the ‘restriction’ of having to initialise the system close to a stationary homogeneous steady state that is unstable in a specific manner, as is the case in the ‘classical Turing’ view of pattern formation, presented at the beginning of section 1.

This does not mean that the process of de novo pattern formation is uninteresting, or not relevant. On the contrary, in many applications, the question how patterns can emerge from a homogeneous background state is central to understanding the underlying biological process. In this context, the novel mechanochemical feedback paradigm offers an attractive and viable alternative to the ‘classical Turing’ theory of de novo pattern formation. This paradigm change also has important consequences for experimental studies of pattern formation: Rather than having to look –perhaps in vain– for a candidate inhibitor, one should pay close attention to the geometrical properties of the surface on which the pattern forms.

The impact of the two viewpoints presented in this paper on questions of pattern observability, experimental verification, and biological practice, will be summarised and discussed in section 4.

## 2 Far-from-equilibrium patterns

From the viewpoint of evolution equations (1.1), a *pattern* can be understood as a (stationary) equilibrium *u*_∗_(*x*) in phase space 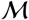, which is spatially non-homogeneous – in particular, which exhibits a clear spatial regularity, e.g. periodicity. Equilibria are, by definition, solutions to (1.3), which is usually called the ‘existence problem’. In the context of reaction-diffusion systems (1.2), this existence problem can take the form of a system of elliptic PDEs (in the case of two or more spatial dimensions), or a system of ODEs (in the case of one spatial dimension). The dimension of *x*, the shape of the spatial domain, the type of boundary conditions, and the specific form of the nonlinearity *F*(*u*) (1.2) all influence the complexity of the existence problem, and the extent to which this problem is analytically tractable.

Once more, dynamical systems theory can, in several important cases, be used to gain insight into pattern solutions. In one spatial dimension, the existence problem (1.3) takes the form of a system of ODEs, in the independent variable *x* ∈ ℝ. This system can now be viewed as a dynamical system in *x*, on the phase space ℝ^2*n*^. For example, if we consider the two-component reaction-diffusion system

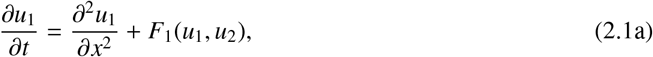

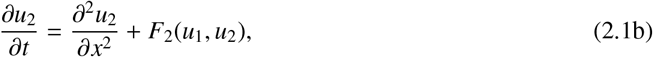

the associated existence problem can be written as the dynamical system

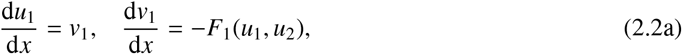

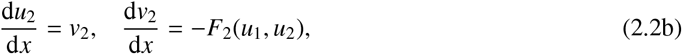

with (*u*_1_, *v*_1_, *u*_2_, *v*_2_) ∈ ℝ^4^. In this viewpoint, patterns are identified as *special orbits* of the dynamical system: Spatially periodic patterns are equivalent to periodic orbits, spatially localised (pulse type) solutions to homoclinic orbits, and wave fronts to heteroclinic orbits. This viewpoint of pattern formation, which is sometimes called *spatial dynamics* [30, 31, 32, 33, 34], aims to use techniques and methods for the analysis of special solutions that have been developed in the context of dynamical systems.

### 2.1 Spatial scale separation and geometric singular perturbation theory

Nevertheless, identifying a suitable mathematical context does not immediately imply that real insight into the existence and shape of patterns can be gained. High-dimensional, nonlinear dynamical systems can –and often will– exhibit chaotic and highly complex behaviour, posing a clear difficulty for the explicit analysis of pattern-like structures.

Here, the biological context offers a way forward, alleviating these mathematical difficulties by introducing *spatial scale separation*. Almost without exception, biological models for pattern formation in a specific context explicitly incorporate the observation that different model components act on different spatial scales. Not only is this spatial scale separation seen as a prerequisite for pattern formation (cf. the LALI principle); in addition, model components are frequently *defined* by the spatial scale on which they act – thereby ingraining this principle of spatial scale separation in the core of the model. In the context of reaction-diffusion equations, this principle is implemented by the assumption that the ratio of the diffusivity rates of two model species is *small*, that is, 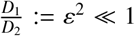. This small parameter ε now introduces a scale separation in the existence problem, in the following way. Take the example system (2.1). Introducing ε^2^ as the diffusivity of *u*_1_ would change the associated existence problem to

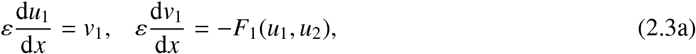

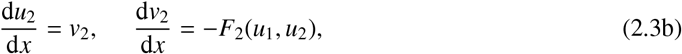

Introducing the short scale variable 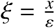, (2.3) transforms to

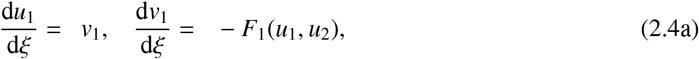

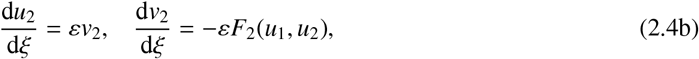

For ε > 0, the dynamics of systems (2.3) and (2.4) are completely equivalent, and their orbit structure is identical. However, their *singular limit* ε → 0 is completely different. System (2.3) reduces to

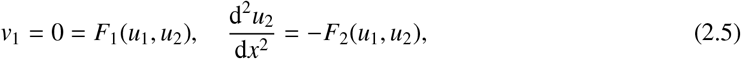

two algebraic equations and a second-order differential equation, while the singular limit of system (2.4) yields

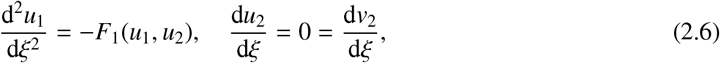

a second-order differential equation and two very simple first-order differential equations. Hence, we can identify (*u*_1_, *v*_1_) as short scale components and (*u*_2_, *v*_2_) as long scale components, presenting a *decomposition* of the original existence problem. This decomposition reduces the model complexity, thereby enabling more detailed and explicit analysis.

The procedure sketched above is made rigorous by *geometric singular perturbation theory* (GSPT), which is based on the pioneering work by Fenichel [35, 36]. GSPT employs a combination of techniques and concepts from geometry and dynamical systems to constructively establish the existence of special orbits, which makes it a perfect candidate for the analysis of pattern formation from the viewpoint of spatial dynamics. By virtue of the geometric techniques, specific properties of the constructed orbit (such as amplitude, shape, spatial period) are found explicitly, as they play a pivotal role in the construction process, which at the same time provides a mathematically rigorous proof of the existence of the pattern. Hence, GSPT not only provides mathematical rigour, but also direct practical applicability of the results for specific models [37]. That is, given a reaction-diffusion system that incorporates spatial scale separation, a successful GSPT analysis will make clear which patterns exist, what these patterns look like, what their period is, and how these properties depend on the model parameters [38, 39].

However, the practical applicability of GSPT transcends specific model contexts. This is because the existence and properties of (special) orbits is directly related to the shape and transverse intersection of certain geometrical objects (stable/unstable manifolds) in phase space, see Figure 3. This has, from a model perspective, two consequences. First, recent results [40, 41] show that the existence (and properties) of several important types of special orbits can be established using only *general* properties of the reaction terms *F*(*u*) (1.2) – that is, patterns such as pulses or periodic orbits, and their properties, can be found for *general classes* of reaction-diffusion systems. For a specific reaction-diffusion system, one only needs to check whether its reaction terms obey certain (mild) conditions (e.g. [40, Assumptions (A1–4)]); if so, the pattern properties are explicitly given in terms of integrals involving the reaction terms, and certain solutions to lower-dimensional differential equations. A second, and related, consequence of the geometric approach, is that because the specific functional form of the reaction terms *F*(*u*) (1.2) is not important, the reaction terms can be directly defined through an (experimentally obtained) response curve. Only the geometric shape of this curve determines the existence and properties of pattern solutions, not the specific algebraic implementation of this shape. Therefore, patterns obtained by a GSPT construction are *structurally stable*. This is particularly important for the connection with biological practice. The fact that mathematical models are, by nature, a strongly reduced idealisation of biological reality, is often a major point of criticism from biologists, who object that the omission of several biologically observed processes that are thought to play a minor role, makes the resulting model less reliable. The geometrical construction provided by GSPT shows that, as long as the resulting response curves do not fundamentally change when these minor processes are included, the mathematical statements regarding patterns remain valid for the extended model.

**Figure 3:**
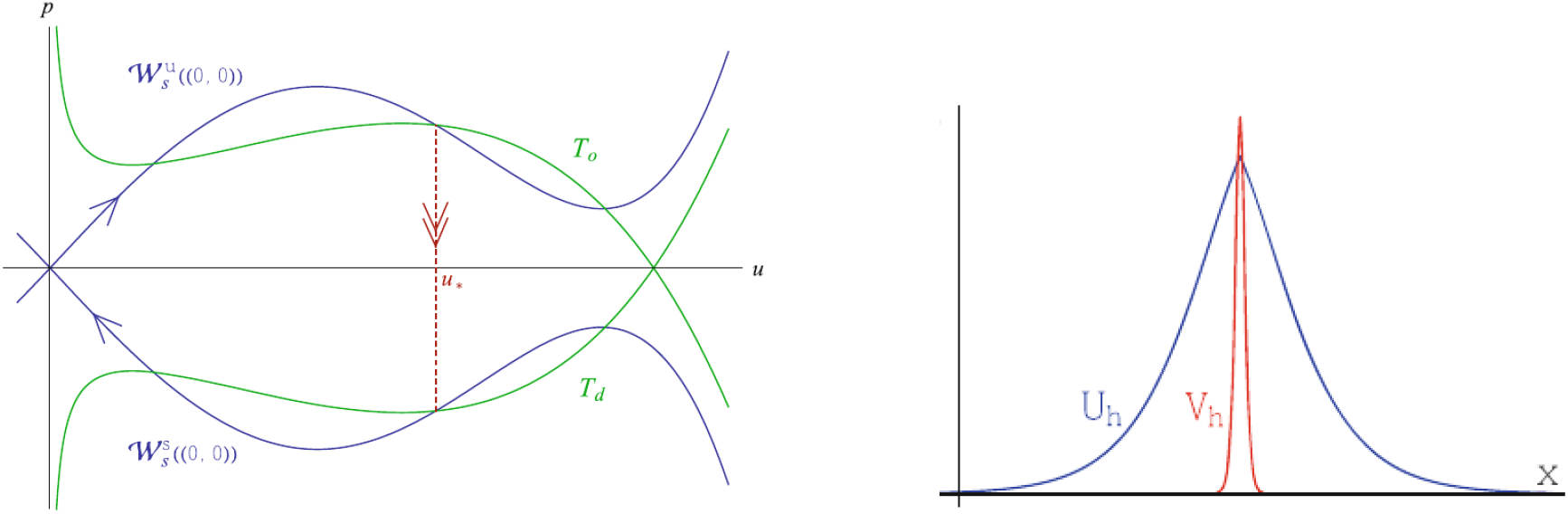
Left: A sketch of the GSPT construction of a homoclinic orbit in a projection of phase space. A homoclinic orbit can be constructed at every point where the blue and green curve have a transverse intersection. Right: The resulting pattern, a localised pulse. Source: [40].

From the mathematical perspective, it is worthwhile to note that the orbits constructed by GSPT are *exact* solutions to the full nonlinear existence problem (1.3), even though their construction uses approximations in the singular limit ε → 0. In contrast to the linear analysis of a Turing bifurcation, the nonlinearity of the reaction terms *F*(*u*) is crucial to the pattern construction procedure. An additional advantage, in comparison with ‘classical’ matched asymptotics approaches, is that one avoids potentially divergent asymptotic series and technicalities such as secular terms; for sufficiently small ε, GSPT rigorously and robustly proves the existence of (singular) orbits.

### 2.2 Pattern stability and the Busse balloon

Solving the existence problem (1.3) by finding a pattern solution is necessary, but not sufficient, to describe the process of pattern formation. In the context of evolution equations (1.1), we have found a candidate stationary state, but it as yet unclear if and when the evolution will steer the system towards this state.

A minimal requirement for a pattern state to be *observable* as a stationary solution to the evolution equation (1.1), is that the pattern needs to be *dynamically stable*. That is, when the stationary pattern state is perturbed, this perturbation should disappear in time, as the system returns to the pattern state. In other words, when the system is initialised at a point in phase space close to the pattern equilibrium, the (local) flow of the evolution equation should point towards the equilibrium. Hence, a stable pattern is a *local attractor* in the phase space of (1.1).

The local behaviour of the flow of (1.1) in the neighbourhood of an equilibrium *u*_∗_ (1.3) can be determined by linearisation. In the context of reaction-diffusion equations, one obtains a linear, parabolic, system of PDEs which explicitly (i.e. non-autonomously) depend on *x*, because we linearise ‘around’ a pattern solution, which has spatial structure – indeed, that last property is at the heart of our endeavour. For the two-component reaction-diffusion system (2.1), the associated linearised system around a given stationary pattern solution (*u*_1_(*x*), *u*_2_(*x*)) would take the form

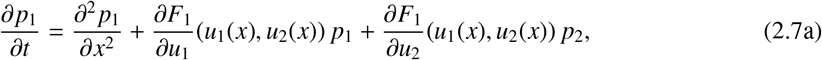

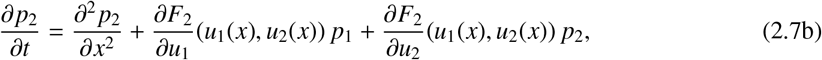

where (*p*_1_(*x*, *t*), *p*_2_(*x*, *t*)) is the perturbation. The pattern (*u*_1_(*x*), *u*_2_(*x*)) is linearly (asymptotically) stable if (*p*_1_(*x*, *t*), *p*_2_(*x*, *t*)) → (0, 0) as *t* → ∞.

Due to the presence of non-autonomous terms, the analysis of systems such as (2.7) is difficult in general, making it challenging to obtain information about the temporal behaviour of perturbations (*p*_1_, *p*_2_). However, when the original system exhibits spatial scale separation, this feature is carried over to the linearised problem. In turns out that, by taking an approach similar to that taken in the existence problem (GSPT), one can obtain detailed information on the temporal behaviour of perturbations, and derive explicit conditions for stability of the underlying pattern [40, 41]. Again, the spatial scale separation makes all the difference: It turns an intractable problem into a manageable one.

Whether one employs analytical techniques or numerical simulations to investigate patterns and their stability: Given a specific biological system, one would like to have a clear overview of the set of stable patterns. The Busse balloon [42, 43, 44] provides such an overview, see Figure 4 A. Here, the focus is on patterns with a clearly defined wavelength, which can therefore be used to characterise the pattern. The Busse ballon is defined as the set in (parameter, wave number) space where patterns with that wave number are dynamically stable for that parameter combination. The specific shape of the Busse balloon can vary significantly from system to system, and the Busse balloon sketched in Figure 4 A captures some important pattern formation phenomena, but certainly not all. Perhaps most importantly, the Busse balloon does not incorporate the amplitude of the pattern, which means close-to-equilibrium and far-from-equilibrium patterns cannot be distinguished in this view. As another example, given a parameter combination for which multiple (sometimes, a range of) stable patterns exist, the Busse balloon does not tell us which pattern is *selected* – that is, which pattern is ‘the most stable’, or which pattern has the largest ‘basin of attraction’ in the phase space of (1.1). The answer to this question may even, for example, depend on the rate of change of the system parameter, see [44, Figure 12].

**Figure 4:**
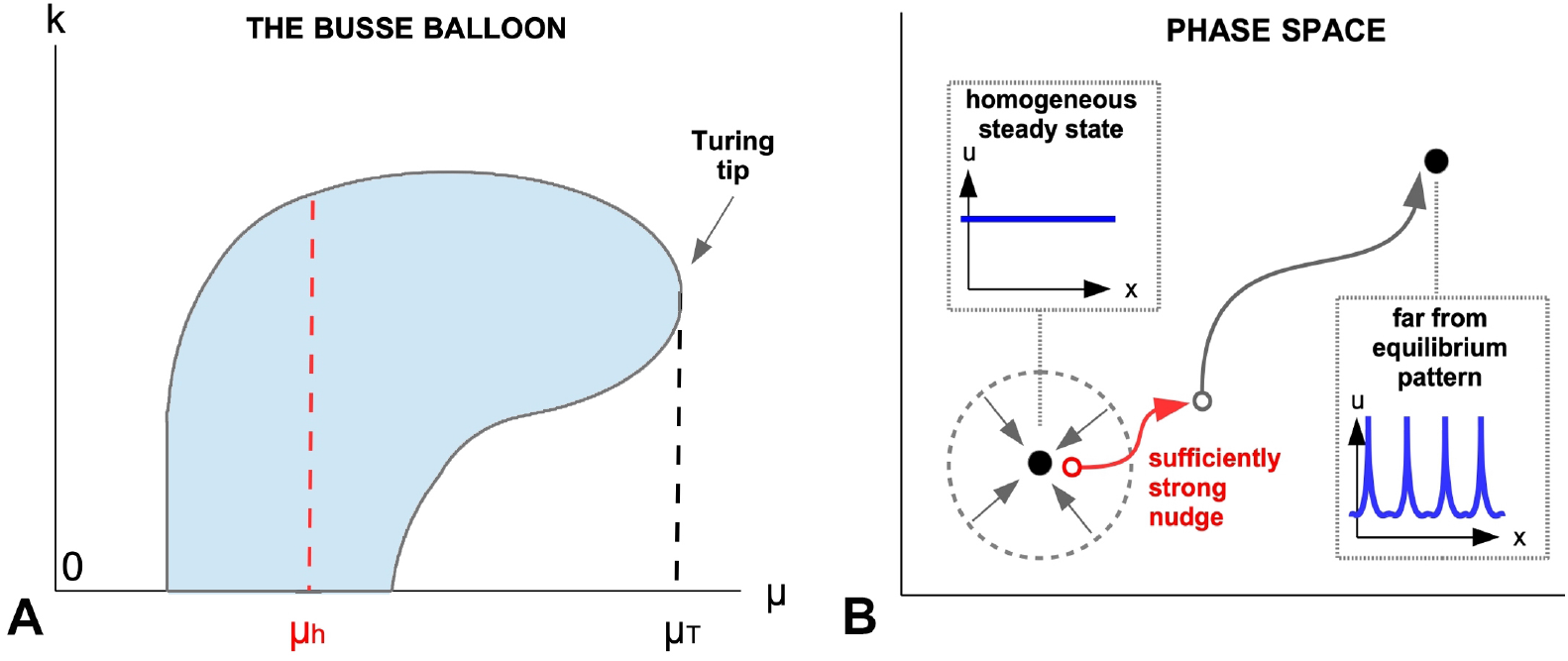
(A) The Busse balloon in (μ, *k*) space, where μ is a generic system parameter, and *k* the wave number; spatially homogeneous steady states have wave number *k* = 0. In the shaded region, patterns with wave number *k* exist and are stable. At the tip of the Busse balloon, a Turing bifurcation takes place for μ = μ_*T*_. For μ < μ_*h*_, the homogeneous background state is stable, yet stable patterns exist. (B) Given a sufficiently strong perturbation (‘nudge’), a system can be forced to evolve towards a far-from-equilibrium pattern, even though the homogeneous stationary state is stable.

With the help of the analytical techniques described in this section, we can investigate the feasibility of the different scenarios identified in section 1.2. For comparison, we add a fourth scenario that incorporates the ‘classical Turing’ view on pattern formation.

1. An initial state *u*_0_ may evolve towards a far-from-equilibrium pattern. The existence of such far-from-equilibrium patterns can be established using GSPT; similar techniques can be used to determine whether an existing pattern is a local attractor. Whether the initial state actually flows towards this stationary pattern solution, depends on the global properties of the evolution equation: The initial state has to lie inside the pattern’s basin of attraction. Note that these considerations do not depend on whether the initial state *u*_0_ is chosen close to a spatially homogeneous steady state; hence, this also incorporates scenario 3.
2. An initial state *u*_0_ may evolve towards a an attractor in phase space that is *not* an equilibrium (1.3), i.e. not a stationary solution, but a fully spatiotemporal solution to (1.1). Establishing the existence of such attractors is a challenging task, but can be be accomplished in the presence of certain system symmetries (e.g. rotating spirals [45]), or in the neighbourhood of bifurcations (e.g. breathing pulses [46]). As in scenario 1, these considerations do not depend on whether the initial state *u*_0_ is chosen close to a spatially homogeneous steady state; hence, this also incorporates scenario 3.
3. The system may evolve towards a stationary pattern, regardless of the proximity of the initial state *u*_0_ to a spatially homogeneous stationary state *u*_∗_; see points 1 and 2. Note that this implies that a stable spatially homogeneous steady state and a stable (far-from-equilibrium) pattern can *coexist*; this phenomenon is also (potentially) present in the Busse balloon shown in Figure 4 for μ < μ_*h*_. From a classical pattern formation point of view, this implies that the system needs a sufficiently large ‘nudge’ to escape the basin of attraction of the homogeneous steady state and evolve towards a pattern. This bistability phenomenon can have profound implications for the understanding of pattern formation in biological systems where a homogeneous steady state is robustly stable.
4. An initial state *u*_0_, that is chosen close to a spatially homogeneous steady state, evolves to a close-to-equilibrium pattern with a small amplitude; this is the ‘classical Turing scenario’. This means that a pattern solution close to the steady state must be shown to exist. The existence and stability of close-to-equilibrium patterns for general systems can be determined using modulation equations and related techniques (e.g. [47, 48, 49]).

## 3 Mechano-chemical feedback: A new paradigm for pattern formation

Recent experimental research indicates that besides molecular signalling, also tissue mechanics play an active role in tissue patterning [50, 51, 52, 53]. Motivated by the lack of experimental identification of the long-range inhibitor [54, 55], the mechano-chemical concept of pattern formation offers a viable alternative to the ‘classical Turing’ theory that may have important consequences for experimental studies. Among others, this yields a question whether the interplay between chemical and mechanical processes matches the LALI principle, e.g., by mechanical cues providing the long-range inhibition.

A first toy model demonstrating pattern formation potential of mechano-chemical feedback loops has been presented in [17]. It is based on existence of one diffusing morphogen that yields tissue evaginations. In turn, the tissue evagination stimulates production of the morphogen. The positive feedback loop results in a local self-activation, accompanied by a long-range inhibition. The latter effect emerges since local outward bending (relative to the initial curvature) always induces inward bending at the boundaries of the evaginated patch. We consider evolution of a closed 2D tissue surface Γ given by a parametric representation 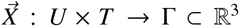 and a morphogen concentration ϕ : *U* → ℝ_≥0_ naturally moving with the deforming tissue (by identifying 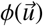 with 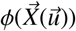. Curvature-dependent elastic properties of the tissue are given by the modified Helfrich energy

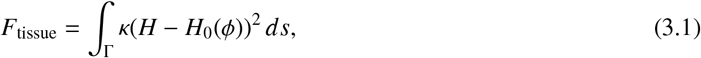

which is an approach frequently used for elastic modelling of thin cell membranes [56, 24, 57, 58]. Here, *H* is the mean curvature, *ds* is the surface measure, the bending rigidity is given by κ, and the preferred local curvature by *H*_0_. *H* is a nonlinear function of the second derivatives of 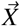; κ and *H*_0_ are elastic parameters representing local mechanical properties. Dependence of *H*_0_ on ϕ reflects the assumption that the morphogen ϕ induces local outward bending (e.g., *H*_0_ ≔ ϕ). The evolution of the tissue 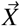 is defined by a *L*^2^–gradient flow of *F*_tissue_ under the constraint of local incompressibility of the tissue. The energy minimisation approach results in a 4th-ordern partial differential equation (PDE):

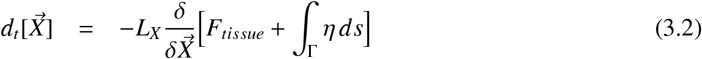

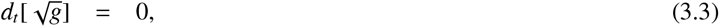

where *d_t_* is the total time derivative, *L_X_* is a kinetic coefficient, 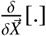 denotes the variation with respect to the arbitrary vector 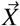, and η is a local Lagrange multiplier [59] due to the constraint of incompressibility, and 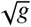 represents the local tissue area (more details are given in [24, 17]).

Morphogen dynamics are given in terms of a reaction-diffusion equation on the deforming surface. Additionally to diffusion and degradation [60, 61, 62], we assume that production of the morphogen ϕ is induced by the local curvature *H* > *H_i_* with initial (mechanically relaxed) curvature 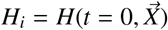. The latter is motivated by some experiments [63, 64, 50]. Using a Michaelis-Menten model for production and defining *H*_≥0_ ≔ max{(*H* − *H_i_*), 0}, we obtain a dynamical equation for ϕ:

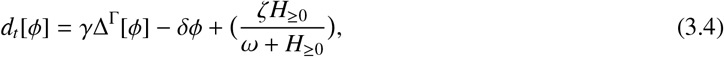

with constants γ, δ, ζ, ω ℝ_≥0_. In summary, the mechano-chemical model is given by a nonlinear 4th-order PDE system, coupling a gradient flow for tissue mechanics (3.2) under the constraint of local area incompressibility (3.3) with a reaction-diffusion equation (3.4) for morphogen dynamics. Model simulations show spontaneous formation of morphogen and curvature patches (Fig. 2 C) that are robust to parameter changes: All examined scenarios led to *de novo* patterns [17]. Further works using 3D continuum mechanical models confirmed that robustness. It was shown that different feedback loops between morphogens and tissue mechanics (measured in terms of strain, stress, or compression) can lead to a spontaneous patterning [65, 27]. Due to the complexity of the structure of the mechano-chemical model, it analysis and numerical implementation are challenging and require application of novel mathematical and numerical techniques.

## 4 Beyond Turing: Bridging the gap between observation and theory

The two approaches presented in this paper represent two distinct views on the study of patterns: pattern *formation* versus pattern *analysis*. Pattern formation is concerned with the initial stages of the pattern forming process, and aims at understanding *de novo* patterns; in this view, the (biological) interpretation of the underlying model and its separate components is central to understanding and interpreting the creation of patterns. In contrast, pattern analysis considers the final stages of a pattern formation process, and aims to study a pattern as a stationary, spatially structured solution to an evolutionary process, which is often modelled using evolution equations. Here, the question is not how a certain pattern arises, but whether it exists and is stable. Both structural and dynamical stability, that is, robustness under both external/random perturbations and internal/systematic perturbations, is necessary for a pattern to be *observable*.

Patterns that are observed in nature are often robust, stationary features. Hence, to understand these patterns in the context of a biological model, the pattern analysis view seems appropriate. After all, ‘classical’ Turing analysis focuses on the emergence of pattern-like structures from a homogeneous steady state, but does not provide any statements about the result of this pattern forming process: This is a clear gap between ‘theoretical understanding’ and ‘experimental observation’. Moreover, not every observed pattern needs to have arisen through a Turing bifurcation, and vice versa, not every Turing bifurcation necessarily leads to an observable, robust, stable pattern.

In addition, the evolution equation view shows that multiple stable patterns can coexist; in particular, a stable homogeneous background state can coexist with a stable pattern, which presents a departure from the ‘classical’ Turing intuition. This phenomenon of bi-or multi-stability can be seen as a sign of robustness of the underlying system, as several steady states can be stable with respect to dynamical or structural perturbations. In contrast, a system that is close to a Turing bifurcation, is very sensitive to perturbations. This induces a degree of unpredictability of the outcome of the system’s evolution: One cannot precisely predict if, and when, a small randomly occurring perturbation may induce spontaneous pattern formation. This lack of control seems to be an undesirable quality of a biological system. For example, the formation of patterns could trigger the next stage of a developmental process, and the timing of this step is crucial for the success of this developmental process. On the other hand, if a system is bi-stable, a sufficiently strong nudge would be needed to perturb the system sufficiently far away from its stable stationary state (see Figure 4 B). The probability of such a strong nudge appearing at random is negligible; therefore, this presents a strong measure of potential control over *de novo* pattern formation, which can be induced only by a targeted, strong, perturbation. In some cases both mechanisms can co-exist resulting in *de novo* initiation of spatially heterogeneous structures that evolve towards *far-from-equilibrium* patterns [22]. Recent simulation studies have demonstrated richness of possible chemical networks that are able to produce patterns following Turing instability [66, 67]. However, simulation results alone do not provide understanding of the nature of the observed patterns. For example, it was shown that in models coupling a reaction-diffusion equation with a system of ordinary differential equations all *close-to-equilibrium* (Turing) patterns are unstable and the solutions that emerge due to Turing instability must be *far-from-equilibrium* patterns [21, 22].

Instead, the viewpoint of pattern formation is much more focused on the details of the biological model, where the creation of *de novo* patterns results from the interplay of different model features, and is seen as a hallmark of, for example, a LALI-type model structure. Hence, a pattern is seen as evidence for LALI, and for the presence of an activator-inhibitor type interaction. However, as the identification of model components that seem to be necessary to produce patterns this way has proven to be difficult, we propose a *paradigm change*. That is, we propose that one should consider the spatial background as an active component in the patterning process. In the most fundamental description, one chemical component diffusing on a surface interacts with that surface through change in curvature, yielding an effective LALI-interaction that produces patterns. This has at least two clear advantages: one does not need to assume the existence of an elusive second chemical component, and one takes the actual shape of the ‘biological’ background into account. More generally, almost all pattern formation analysis thus far has assumed a fixed, flat background, based on the assumption that the shape of the background does not influence the behaviour of the pattern (see [68, 69, 70] for notable exceptions). In the mechano-chemical approach as presented in section 3, we abandon this assumption, and investigate what happens if the background shape *does* matter – not only that, but what if the background plays an active role? In many developmental applications, one observes that the shape of the organism changes according to the pattern that occurs [71]. Until now, this has been considered a response to a previously established pattern. We propose that these two phenomena, i.e. the formation of a pattern and the change of background curvature, occur simultaneously. Moreover, we propose that these two dynamical processes interact, and that the combination of pattern and curvature arises precisely due to this interaction.

Regardless whether one takes the viewpoint of pattern formation or pattern analysis, observed patterns are regarded as large-scale features that arise from small-scale, sometimes microscopic, interactions. As such, patterns as emergent phenomena can be used to make complex microscopic interactions experimentally accessible. When one adopts this line of reasoning, two questions present themselves. First: are the ‘canonical’ models that are widely in use, able to produce all types of patterns that have been observed? Second: given an observed pattern, is it possible to infer the structure of the underlying model? The answer to the first question highly depends on what one considers to be ‘canonical’. However, recent pattern analysis research has shown that well-known reaction-diffusion models such as the Gierer-Meinhardt, Gray-Scott/Klausmeier or Schnakenberg model only admit a restricted class of far-from-equilibrium patterns – that is, when one considers reaction-diffusion systems with sufficiently general nonlinear reaction terms, one suddenly has access to a vastly extended library of pattern shapes. Even more so: for a reaction-diffusion system exhibiting spatial scale separation, thereby allowing the application of GSPT, one can let the pattern shape *define* the nonlinear reaction terms in the model. Moreover, as the temporal behaviour of far-from-equilibrium patterns in general reaction-diffusion systems strongly depends on the specific shape of the reaction terms, the stability [72], the spatio-temporal dynamics of patterns [46], and even the shape of the Busse balloon [73] can be tuned, to yield novel pattern dynamics. The second question is of fundamental importance when one aims to use observed patterns for model comparison purposes. Not only is this closely related to the question to which degree the presence of hypothesised microscopic processes can be experimentally verified; it also directly impacts model benchmarking, model selection, and parameter identifiability. Recent research has shown that the relation between observed patterns and candidates for the underlying model is far from one-to-one: On one hand, an observed pattern can be matched to several distinct pattern generating models; on the other hand, patterns that look identical to the naked eye can be distinguished using statistical measures, and can hence be shown to originate from different models [74].

## Acknowledgements

This work is supported by Deutsche Forschungsgemeinschaft (DFG) under Germany’s Excellence Strategy EXC-2181/1 - 390900948 (the Heidelberg STRUCTURES Excellence Cluster) and SFB1324 (B05). FV was supported by a Humboldt Research Fellowship.

## References

[1] A. Gierer and H. Meinhardt, “A theory of biological pattern formation.” Kybernetik, vol. 12, no. 1, pp. 30–9, 1972.

[2] A. M. Turing, “The chemical basis of morphogenesis.” Phil. Trans. R. Soc. Lond. B, vol. 237, pp. 37–72, 1953.

[3] H. Meinhardt, “A model for pattern formation of hypostome, tentacles, and foot in Hydra: how to form structures close to each other, how to form them at a distance,” Dev. Biol., vol. 157, no. 2, pp. 321–33, 1993.

[4] H. Meinhardt and A. Gierer, “Pattern formation by local self-activation and lateral inhibition.” Bioessays, vol. 22, pp. 753–60, 2000. [Online]. Available: http://dx.doi.org/3.0.CO;2-Z

[5] H. Meinhardt, “Turing’s theory of morphogenesis of 1952 and the subsequent discovery of the crucial role of local self-enhancement and long-range inhibition,” Interface Focus, vol. 2, pp. 407–16, 2012.

[6] S. Kondo and R. Asai, “A reaction–diffusion wave on the skin of the marine angelfish pomacanthus,” Nature, vol. 376, p. 765–768, 1995.

[7] G. von Dassow, E. Meir, E. M. Munro, and G. Odell, “The segment polarity network is a robust developmental module,” Nature, vol. 406, pp. 188–192, 1995.

[8] A. Nakamasu, G. Takahashi, A. Kanbe, and S. Kondo, “Interactions between zebrafish pigment cells responsible for the generation of Turing patterns.” Proc. Natl. Acad. Sci. U S A, vol. 106, no. 21, pp. 8429–34, May 2009.

[9] P. K. Maini, R. E. Baker, and C.-M. Chuong, “Developmental biology. the Turing model comes of molecular age.” Science (New York, N.Y.), vol. 314, pp. 1397–1398, Dec. 2006.

[10] K. Tori, “Two-dimensional spatial patterning in developmental systems,” Trends in Cell Biology, vol. 22, pp. 438–446, 2010.

[11] L. Morelli, K. Uriu, S. Ares, and A. Oates, “Computational approaches to developmental patterning,” Science, vol. 336, pp. 187–191, 2012.

[12] E. Keller and L. Segel, “A model of chemotxis,” J. Theor. Biol., vol. 30, p. 225–234, 1971.

[13] T. Hillen and K. J. Painter, “A user’s guide to pde models for chemotaxis,” J. Math. Biol., vol. 58, p. 183, 2008.

[14] A. Anderson, M. Chaplain, E. Newman, R. Steele, and A. Thompson, “Mathematical modelling of tumour invasion and metastasis,” J. Theor. Med., vol. 2, pp. 129–154., 2000.

[15] B. C. Goodwin and M. H. Cohen, “A phase-shift model for the spatial and temporal organization of developing systems.” J. Theor. Biol., vol. 25, pp. 49–107, Oct. 1969.

[16] J. Murray and G. Oster, “Cell traction models for generating pattern and form in morphogenesis,” J. Math. Biol., vol. 19, pp. 265–279., 1984.

[17] M. Mercker, D. Hartmann, and A. Marciniak-Czochra, “A mechanochemical model for embryonic pattern formation: coupling tissue mechanics and morphogen expression.” PLoS One, vol. 8, no. 12, p. e82617, 2013.

[18] A. Gierer, S. Berking, H. Bode, C. David, K. Flick, G. Hansmann, H. Schaller, and E. Trenkner, “Regeneration of Hydra from reaggregated cells,” Nature New Biol., vol. 239: 91, pp. 98-101, 1972.

[19] S. Kondo, “An updated kernel-based Turing model for studying the mechanisms of biological pattern formation,” J. Theor. Biology, vol. 414, p. 120–1127, 2017.

[20] A. Marciniak-Czochra, S. Härting, G. Karch, and K. Suzuki, “Dynamical spike solutions in a nonlocal model of pattern formation,” Nonlinearity., vol. 31, p. 1757, 2018.

[21] A. Marciniak-Czochra, G. Karch, and K. Suzuki, “Instability of Turing patterns in reaction-diffusion-ODE systems,” J. Math. Biol., vol. 74, pp. 583–618, 2017.

[22] S. Härting, A. Marciniak-Czochra, and I. Takagi, “Stable patterns with jump discontinuity in systems with Turing instability and hysteresis,” Disc. Cont. Dyn. Syst. A., vol. 37, pp. 757–800, 2017.

[23] A. Köthe, A. Marciniak-Czochra, and I. Takagi, “Hysteresis-driven mechanism of pattern formation in a basic reaction-diffusion-ode model,” Disc. Cont. Dyn. Systems-A, vol. 40, p. 3595, 2020.

[24] M. Mercker, A. Marciniak-Czochra, T. Richter, and D. Hartmann, “Modeling and computing of deformation dynamics of inhomogeneous biological surfaces,” SIAM Journal on Applied Mathematics, vol. 73(5), pp. 1768–1792, 2013.

[25] H. Meinhardt, “Modeling pattern formation in hydra: A route to understanding essential steps in development.” Int. J. Dev. Biol., vol. 56, no. 6-8, pp. 447–62, 2012.

[26] M. C. Vogg, L. Beccari, L. Iglesias Ollé, C. Rampon, S. Vriz, C. Perruchoud, Y. Wenger, and B. Galliot, “An evolutionarily-conserved Wnt3/β-catenin/Sp5 feedback loop restricts head organizer activity in Hydra,” Nat. Commun., vol. 10, p. 312, Jan. 2019.

[27] F. Brinkmann, M. Mercker, T. Richter, and A. Marciniak-Czochra, “Post-Turing tissue pattern formation: Advent of mechanochemistry.” PLoS Comput. Biol., vol. 14, p. e1006259, 2018.

[28] B. Sandstede, A. Scheel, and C. Wulff, “Bifurcations and dynamics of spiral waves,” Journal of Nonlinear Science, vol. 9, pp. 439 – 478, 1999.

[29] B. Fiedler and A. Scheel, “Spatio-temporal dynamics of reaction-diffusion patterns,” in Trends in Nonlinear Analysis, M. Kirkilionis, S. Krömker, R. Rannacher, and F. Tomi, Eds. Springer Berlin Heidelberg, 2003, pp. 23 – 152.

[30] M. Beck, “Spectral stability and spatial dynamics in partial differential equations,” Notices of the American Mathematical Society, vol. 67, no. 4, pp. 500 – 507, 2020.

[31] K. Kirchgässner, “Wave-solutions o reversible systems and applications,” Journal of Differential Equations, vol. 45, no. 1, pp. 113 – 127, 1982.

[32] A. Mielke, “A reduction principle for nonautonomous systems in infinte-dimensional spaces,” Journal of Differential Equations, vol. 65, no. 1, pp. 68 – 88, 1986.

[33] D. Peterhof, B. Sandstede, and A. Scheel, “Exponential dichotomies for solitary-wave solutions of semilinear elliptic equations on infinite cylinders,” Journal of Differential Equations, vol. 140, no. 2, pp. 266 – 308, 1997.

[34] M. Beck, G. Cox, C. Jones, Y. Latushkin, and A. Sukhtayev, “A dynamical approach to semilinear elliptic equations,” Annales de l’Institut Henri Poincaré C, Analyse non linéaire, vol. 38, no. 2, pp. 421 – 450, 2021.

[35] N. Fenichel, “Persistence and smoothness of invariant manifolds for flows,” Indiana University Mathematics Journal, vol. 21, pp. 193 – 226, 1971.

[36] N. Fenichel, “Geometric singular perturbation theory for ordinary differential equations,” Journal of Differential Equations, vol. 31, no. 1, pp. 53 – 98, 1979.

[37] G. Hek, “Geometric singular perturbation theory in biological practice,” Journal of Mathematical Biology, vol. 60, no. 3, pp. 347 – 386, 2010.

[38] A. Doelman, R. Gardner, and T. Kaper, “Large stable pulse solutions in reaction-diffusion equations,” Indiana University Mathematics Journal, vol. 50, no. 1, pp. 443–507, 2001.

[39] A. Doelman, T. Kaper, and H. van der Ploeg, “Spatially periodic and aperiodic multi-pulse patterns in the one-dimensional Gierer-Meinhardt equation,” Methods and Applications of Analysis, vol. 8, no. 2, pp. 387–414, 2001.

[40] A. Doelman and F. Veerman, “An explicit theory for pulses in two component, singularly perturbed, reaction-diffusion equations,” Journal of Dynamics and Differential Equations, vol. 27, no. 3, pp. 555–95, 2015.

[41] B. de Rijk, A. Doelman, and J. D. M. Rademacher, “Spectra and stability of spatially periodic pulse patterns: Evans function factorization via Riccati transformation,” SIAM Journal on Mathematical Analysis, vol. 48, no. 1, pp. 61–121, 2016.

[42] F. Busse, “Non-linear properties of thermal convection,” Rep. Prog. Phys., vol. 41, no. 12, pp. 1929– 67, 1978.

[43] S. van der Stelt, A. Doelman, G. Hek, and J. Rademacher, “Rise and fall of periodic patterns for a generalized Klausmeier-Gray-Scott model,” Journal of Nonlinear Science, vol. 23, no. 1, pp. 39 – 95, 2013.

[44] E. Siero, A. Doelman, M. Eppinga, J. Rademacher, M. Rietkerk, and K. Siteur, “Striped pattern selectio by advective reaction-diffusion systems: Resilience of banded vegetation on slopes,” Chaos, vol. 25, p. 036411, 2015.

[45] B. Sandstede and A. Scheel, “Spiral waves: linear and nonlinear theory,” Memoirs of the Americal Mathematical Society, 2021. [Online]. Available: https://arxiv.org/abs/2002.10352

[46] F. Veerman, “Breathing pulses in singularly perturbed reaction-diffusion systems,” Nonlinearity, vol. 28, no. 7, pp. 2211–46, 2015.

[47] P. Kirrmann, G. Schneider, and A. Mielke, “The validity of modulation equations for extended systems with cubic nonlinearities,” Procedings of the Royal Society of Edinburgh Section A: Mathematics, vol. 122, no. 1-2, pp. 85–91, 1992.

[48] A. Mielke and G. Schneider, “Attractors for modulation equations on unbounded domains – existence and comparison,” Nonlinearity, vol. 8, no. 5, pp. 743–768, 1995.

[49] J. Guillod, G. Schneider, P. Wittwer, and D. Zimmermann, “Nonlinear stability at the Eckhaus boundary,” SIAM Journal on Mathematical Analysis, vol. 50, no. 5, pp. 4699–4720, 2018.

[50] E. Farge, “Mechanotransduction in development.” Curr Top Dev Biol, vol. 95, pp. 243–265, 2011. [Online]. Available: http://dx.doi.org/10.1016/B978-0-12-385065-2.00008-6

[51] S. A. Braybrook and A. Peaucelle, “Mechano-chemical aspects of organ formation in *Arabidopsis thaliana*: the relationship between auxin and pectin.” PLoS One, vol. 8, no. 3, p. e57813, 2013.

[52] E. Hannezo and C.-P. Heisenberg, “Mechanochemical feedback loops in development and disease.” Cell, vol. 178, pp. 12–25, Jun. 2019.

[53] E. Braun and K. Keren, “Hydra regeneration: Closing the loop with mechanical processes in morphogenesis,” BioEssays, vol. 40, p. 1700204, 2018.

[54] S. Kondo and T. Miura, “Reaction-diffusion model as a framework for understanding biological pattern formation.” Science, vol. 329, no. 5999, pp. 1616–20, Sep 2010.

[55] T. W. Hiscock and S. G. Megason, “Mathematically guided approaches to distinguish models of periodic patterning.” Development, vol. 142, no. 3, pp. 409–19, Feb 2015.

[56] M. Mercker and A. Marciniak-Czochra, “Bud-neck scaffolding as a possible driving force in escrt-induced membrane budding.” Biophys J, vol. 108, no. 4, pp. 833–843, Feb 2015.

[57] M. Mercker, M. Ptashnyk, J. Kühnle, D. Hartmann, M. Weiss, and W. Jäger, “A multiscale approach to curvature modulated sorting in biological membranes,” J Theo Biol, vol. 301, pp. 67–82, 2012.

[58] M. Mercker, T. Richter, and D. Hartmann, “Sorting mechanisms and communication in phase-separating coupled monolayers.” J Phys Chem B, vol. 115, pp. 11 739–11 745, 2011.

[59] G. Foltin, “Dynamics of incompressible fluid membranes.” Phys Rev E Stat Phys Plasmas Fluids Relat Interdiscip Topics, vol. 49, pp. 5243–5248, 1994.

[60] B. Alberts, D. Bray, and J. Lewis, Molecular biology of the cell. Garland Publishing, Inc., 2006.

[61] T. Gregor, E. F. Wieschaus, A. P. McGregor, W. Bialek, and D. W. Tank, “Stability and nuclear dynamics of the bicoid morphogen gradient.” Cell, vol. 130, pp. 141–152, 2007.

[62] I. The and N. Perrimon, “Morphogen diffusion: the case of the wingless protein.” Nat Cell Biol, vol. 2, pp. E79–E82, 2000.

[63] E. Brouzes and E. Farge, “Interplay of mechanical deformation and patterned gene expression in developing embryos.” Curr. Opin. Genet. Dev., vol. 14, pp. 367–74, 2004.

[64] N. Desprat, W. Supatto, P.-A. Pouille, E. Beaurepaire, and E. Farge, “Tissue deformation modulates twist expression to determine anterior midgut differentiation in *Drosophila* embryos.” Dev Cell, vol. 15, pp. 470–477, 2008.

[65] M. Mercker, F. Brinkmann, A. Marciniak-Czochra, and T. Richter, “Beyond Turing: mechanochemical pattern formation in biological tissues.” Biol. Direct, vol. 11, no. 22, pp. 1–15, May 2016.

[66] N. S. Scholes, D. Schnoerr, M. Isalan, and M. P. Stumpf, “A comprehensive network atlas reveals that Turing patterns are common but not robust,” Cell Sys., vol. 9, no. 3, pp. 243–57, 2019.

[67] X. Diego, L. Marcon, P. Müller, and J. Sharpe, “Key features of Turing systems are determined purely by network topology,” Phys. Rev. X, vol. 8, no. 2, p. 021071, 2018.

[68] P. Trinh and M. Ward, “The dynamics of localized spot patterns for reaction-diffusion systems on the sphere,” Nonlinearity, vol. 29, no. 3, pp. 766–806, 2016.

[69] J. Tzou and L. Tzou, “Spot patterns of the Schnakenberg reaction-diffusion system on a curved torus,” Nonlinearity, vol. 33, no. 2, pp. 643–674, 2019.

[70] R. Van Gorder, V. Klika, and A. Krause, “Turing conditions for pattern forming systems on evolving manifolds,” Journal of Mathematical Biology, vol. 82, no. 4, pp. 1 – 61, 2021.

[71] S. Gilbert, Developmental Biology. Sinauer Associates, Inc.; 10 edition, 2013.

[72] F. Veerman and A. Doelman, “Pulses in a Gierer-Meinhardt equation with a slow nonlinearity,” SIAM Journal on Applied Dynamical Systems, vol. 12, no. 1, pp. 28 – 60, 2013.

[73] A. Doelman, J. Rademacher, B. de Rijk, and F. Veerman, “Destabilization mechanisms of periodic pulse patterns near a homoclinic limit,” SIAM Journal on Applied Dynamical Systems, vol. 17, no. 2, pp. 1833–90, 2018.

[74] A. Kazarnikov and H. Haario, “Statistical approach for parameter identification by Turing patterns,” Journal of Theoretical Biology, vol. 501, p. 110319, 2020.

